# YACHT: an ANI-based statistical test to detect microbial presence/absence in a metagenomic sample

**DOI:** 10.1101/2023.04.18.537298

**Authors:** David Koslicki, Stephen White, Chunyu Ma, Alexei Novikov

## Abstract

In metagenomics, the study of environmentally associated microbial communities from their sampled DNA, one of the most fundamental computational tasks is that of determining which genomes from a reference database are present or absent in a given sample metagenome. While tools exist to answer this question, all existing approaches to date return point estimates, with no associated confidence or uncertainty associated with it. This has led to practitioners experiencing difficulty when interpreting the results from these tools, particularly for low abundance organisms as these often reside in the “noisy tail” of incorrect predictions. Furthermore, no tools to date account for the fact that reference databases are often incomplete and rarely, if ever, contain exact replicas of genomes present in an environmentally derived metagenome. In this work, we present solutions for these issues by introducing the algorithm `YACHT`: **Y**es/No **A**nswers to **C**ommunity membership via **H**ypothesis **T**esting. This approach introduces a statistical framework that accounts for sequence divergence between the reference and sample genomes, in terms of average nucleotide identity, as well as incomplete sequencing depth, thus providing a hypothesis test for determining the presence or absence of a reference genome in a sample. After introducing our approach, we quantify its statistical power as well as quantify theoretically how this changes with varying parameters. Subsequently, we perform extensive experiments using both simulated and real data to confirm the accuracy and scalability of this approach. Code implementing this approach, as well as all experiments performed, is available at https://github.com/KoslickiLab/YACHT.

## 1 Introduction

### 1.1 Background and context

In the field of metagenomics, that is, the study of microbial communities via their environmentally sampled DNA, there has been increasing interest in the so-called “rare microbiome” [1,2,3,4,5,6]. These are the microbial organisms or taxa at low relative abundance in a given sample where many ecologically important microbes are found [7]. However, it has been widely recognized that computational approaches to inferring the presence and relative abundance of microbes or taxa in given metagenomic sample, so called “profiling techniques” often suffer from the inability to distinguish between statistical noise and microbes at low relative abundance [8,9,10,11]. As a result, particularly in taxonomic profiling, practitioners often resort to filtering their profiles based on some abundance threshold: ignoring tool predictions of taxa with relative abundance below some threshold [12], or else removing the lowest abundance predictions until some fixed amount of abundance has been removed [11]. Such approaches are ad-hoc, with little guidance in how to choose the thresholds [2] and have demonstrated negative effects on community analysis and interpretation [13,14] as well as on profiling tool performance [11]. While some more sophisticated approaches to filtering exist, these are usually limited to specific numeric quantities of interest (such as preserving mutual information [15] or maximizing covariance [16]) or else ignore biologically relevant considerations such as molecular evolution, sequencing error, and low sequencing sampling rates. As such, current microbiome practitioners have few biologically interpretable tools at their disposal to confidently claim a particular microbe is present at low abundance in their sample, or else reject it as noise. Clearly, this has negative impacts on the study of the rare microbiome and has even resulted in high-profile cases of misinterpreting the presence or absence of microbes in a sample (eg. Plague in the New York subway system [17,18,19]).

In this manuscript, we propose to address this situation by developing a theoretically sound statistical test for the presence or absence of a microbe in a metagenomic sample while accounting for sequencing depth constraints and evolutionary divergence between reference and sample genomes. After describing the method and proving its soundness, we demonstrate the effectiveness of this approach with thorough and realistic *in silico* “spike-in” [20] experiments.

### 1.2 Summary of approach

Before describing our proposed approach rigorously, we provide a high-level idea of the approach and its biological interpretation. Two recent innovations have allowed for this approach to be developed: a) a statistical framework in which to study sequence evolution via alignment-free, *k*-mer-based approaches [21], b) a novel data reduction technique [22] and associated metagenomic profiler [23]. The former allows us to relate *k*-mer-based statistics (such as the Jaccard or containment index) to biologically relevant quantities such as Average Nucleotide Identity (ANI) [24,25] while also providing associated confidence intervals and hypothesis tests. The latter innovation provides us a means to quickly derive the prerequisite *k*-mer-based quantities while also providing a theoretical framework in which hypothesis tests can be developed (as we do below) that, among other things, account for sequencing depth variability.

Our approach requires the following information from a user: first, a user specifies a desired false negative rate *α* which controls for the sensitivity of the method in predicting microbial presence or absence. Next the user indicates at what value *A* of Average Nucleotide Identity they consider two organisms to be biologically identical. Using this ANI threshold allows us to account for evolutionary sequence divergence between reference and sample genomes, as well as account for sequencing error. Lastly, the user indicates what fraction *c* of the genetic material exclusive to a given microbe’s genome needs to be present in the sample (i.e. its coverage [26]) for the microbe itself to be called “in the sample”. As input, the method takes a reference database of microbial genomes and an unprocessed metagenomic sequencing sample.

From here, the method then identifies organisms in the reference database that could possibly be contained in the sample: those with non-zero *k*-mer overlap for a large value of *k* (eg. *k* = 31, 51, etc.). Among these candidate microbes, the method then identifies which *k*-mers are exclusive to each genome. With these in hand, a hypothesis test is performed for each candidate genome that answers the question:

Does the sample contain enough of the *k*-mers exclusive to this reference genome to support the claim that an organism, with ANI at least *A* to this reference genome, is contained in the sample with coverage at least *c*?

If the answer is in the affirmative, then this reference organism is said to be *present* in the sample. Importantly, this process is conducted in an alignment-free fashion, facilitating large-scale analyses and ever expanding reference databases.

### 1.3 Contribution

To our understanding, this is the first approach that disentangles the difference between low-abundance sample organisms and those that are diverged from a given reference database. Due to sampling depth constraints, low-abundance organisms present in a sample often have only a fraction of their genome covered by sample reads. Our approach’s ANI threshold *A* and coverage parameter *c* take this into account. Hence, as long as *c* percentage of a genome’s unique *k*-mers are covered by at least one read, confidence can be had as to this genome’s presence or absence in the sample. Experiments demonstrate that *c* can reliably be set lower than 10%, which translates to relative abundances as low as 0.0035%, thus facilitating “rare microbiome” analyses.

A drawback of this method is that genomic evolution and sequencing error cannot be disentangled. Indeed, from a *k*-mer perspective and the mutation model we assume, there is no difference between “the similarity of one genome and a 1 − *A*-mutated version of it”, and “the similarity of one genome and a 1x-coverage sampled version of it that has a sequencing error rate of 1 − *A*”. Fortunately though, *k*-mers that have been created due to sequencing error do not factor into the determination of presence or absence of organisms in a sample for sufficiently large *k*, since only unique *k*-mers matching to the reference are considered in the hypothesis test.

Additionally, this approach addresses the issue with incomplete reference databases which has been observed to impede metagenomic profiling and analysis approaches [27,28,29,30]. It achieves this since only a mutated version of a sample organism within the ANI threshold need be present in a reference database for detection to be possible.

An additional drawback of this approach is that, in contrast to clade-specific marker gene approaches [31,32,33] or clade-specific *k*-mer based approaches [34,35], our method, when used as a taxonomic profiler, does not attempt to classifier to higher, internal taxonomic ranks. This is due to the fact that average nucleotide (or amino acid) identity does not harmonize with taxonomy, though there have been efforts to re-define taxonomy to address this discrepancy [36,37]. As such, we deemphasize ascertaining taxonomic similarity and instead focus on ANI similarity.

Of note, the framework we have developed is applicable to more than just the detection of organisms in a metagenomic sample. Indeed, our method requires only a reference database and a sample consisting of reads from a portion of the database which have undergone mutations and/or were incompletely sampled. Hence, it is straightforward to extend this approach to applications such as functional annotation (where the reference consists of genes or protein families, and the sample is inferred protein coding sequences), metatranscriptomics (where the reference consists of transcripts and the sample is a shotgun RNA-seq sample), etc.

## 2 Algorithm

In this section, we introduce the YACHT algorithm for detecting organisms in a metagenomic sample and the associated mathematical model.

### 2.1 Preliminaries

We now place the problem of metagenomic presence/absence detection in a mathematical setting. We begin by establishing the following parameters for our model:

- *k* ∈ ℕ, the *k*-mer size,
- *K* ∈ ℕ, the total number of *k*-mers sampled across all organisms,
- *N*, the number of known reference genomes,
- *A* ∈ (0, 1), the ANI threshold above which organisms are said to be distinct.

The intuition and parameter tuning behind the YACHT algorithm are based on the simple mutation model outlined in [21]. In this model, given a specified ANI *a* and string of nucleotides *S* (the genome), a mutation occurs independently at each site with probability 1 − *a*, in which case that nucleotide is changed uniformly at random into one of the other three.

We assume knowledge of a reference database 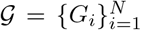 of genomes’ *k*-mers, where each *G*_*i*_ is the set of all *k*-mers of the *i*-th reference genome. In practice, computational necessity means we observe only a “sketch” consisting of subsets of each *G*_*i*_. By a minor abuse of notation, we will use *G*_*i*_ to represent the set of sketched *k*-mers. We use FracMinHash [23,22] as the sketching approach since it has the desirable feature that if a *k*-mer appears in two genomes *i* and *j* and is in the sketch *G*_*i*_ then it will also be the sketch *G*_*j*_. FracMinHash has a “scale factor” *s* which represents the fraction of *k*-mers that are sampled by the sketch, hence 0 *< s* ≤ 1. A FracMinHash **sketch** of *k*-mers is a uniformly random selection of *s*% of those *k*-mers. This is achieved by taking a fixed, uniform hash function *h* with a domain of all *k*-mers and range [0, 4^*k*^] and selecting those *k*-mers *x* such that *h*(*x*) ≤ *s*4^*k*^.

### 2.2 Mathematical model for the sample 𝒮

We will assume that mutations at any site in an organism’s genome are independent from all other mutations in that organism’s genome as well as from mutations in other organism’s genomes. Under this assumption, so long as the scale factor *s* is relatively small and *k* is relatively large, we have the following consequences, which hold with high enough probability that any deviations can be treated as tolerable noise:

- **Uniqueness:** Each *k*-mer appears at most once in any genome.
- **Non-overlap:** Sketched *k*-mers in the reference do not overlap in the genome from which they were derived. Accordingly, the event of a mutation in each sketched *k*-mer will be independent.
- **One-way Mutation:** If a *k*-mer is not in the sketch, the mutation process does not mutate it into a *k*-mer that is in the sketch. Therefore mutation causes *k*-mers to drop out of the sketch, but not to enter it.

Given a genome sketch *G*, then, we can model the sketch *G*^*a*^ of a *mutated copy* of *G* with ANI *a* to the original genome as follows: for each *k*-mer *g* ∈ *G*_*i*_, mutate each site with probability 1 − *a*, which means each *k*-mer will be mutated with probability 1 − *a*^*k*^. By non-overlap, these mutations will be independent. If the *k*-mer *g* is mutated, by one-way mutation, the mutated version will not be in the sketch; moreover, by uniqueness this was the only instance of *k*-mer *g* in the genome represented by *G* and so *g* does not appear in the resulting mutated sketch. Thus the sketch *G*^*a*^ of the mutated genome can be characterized by the process: for each *g* ∈ *G*, mutate it with probability 1 − *a*^*k*^. Then *G*^*a*^ consists of the set of all *k*-mers which survive this process, i.e. which are *not* mutated.

Under this definition, the sketch of a random metagenomic sample can be modeled as the union of many such sketches of randomly mutated copies of known genomes. We now make these definitions precise:

#### Definition 1 (Randomly Mutated Sketch)

*Given k-mer size k, ANI a* ∈ [0, 1], *and finite set of k-mers G, let* {*χ*_*g*_}_*g∈G*_ *be a set of independent Bernoulli random variables taking values in* {0, 1} *with* ℙ (*χ*_*g*_ = 1) = *a*^*k*^. *We define the* ***sketch of a randomly*** *a****-mutated copy of*** *G as the random set G*^*a*^ = {*g* ∈ *G*|*χ*_*g*_ = 1}.

In words, *G*^*a*^ is the set of unmutated *k*-mers in *G* after ungoing the simple mutation process with *ANI a*. Accordingly, if *G* is a sketch of *k*-mers from a genome, then *G*^*a*^ represents the sketch of *k*-mers from a mutated version of that genome with expected ANI *a* to the unmutated version. Similarly, we posit the following mathematical model for the sample 𝒮:

#### Definition 2 (Randomly Mutated Sample Sketch)

*Given k-mer size k and an N -element set of ANI’s* **a**, *let* 𝒢 *be a collection of N finite sets of k-mers G*_*i*_. *Then the* ***sketch of a randomly* a*-mutated sample*** 𝒮^a^ *is defined as the union of a*_*i*_*-mutated copies of G*_*i*_:

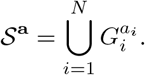

In short, given a set of ANI’s **a**, each genome’s set of *k*-mers *G*_*i*_ ∈ 𝒢 is mutated according to the ANI *a*_*i*_ into a new sketch 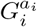. This definition accounts for the case that only a subset of genomes in G are in the sample by setting *a*_*i*_ = 0 for any genomes which are not in the sample, as these will never have any *k*-mers appearing in 𝒮^**a**^ with this model.

The problem of organism membership detection in a sample community can thus be modeled as follows: given a pre-selected ANI threshold *α*, reference sketch 𝒢, and random sample 𝒮^**a**^, determine for each *i* whether a mutated version of genome *i* is in the sample with ANI at least equal to the given threshold: *a*_*i*_ ≥ *A*.

### 2.3 The YACHT algorithm: *N* = 1

For intuition, we first consider the case where our reference G consists of only a single genome sketch *G* containing *K k*-mers. In this case, under the random sample model in Definition 2, our sample sketch S is equal to the sketch of mutated copy of *G* with an unknown ANI *a*, that is: 𝒮^a^ = 𝒢^a^.

Since all *k*-mers in each genome are assumed to have the same mutation probability, the information in the set 𝒮^*a*^ is equivalent to its size |S^*a*^|. Under the model of Definition 2 with *N* = 1, |𝒮^*a*^| = |𝒢^*a*^| = ∑_*g∈G*_ *χ*_*g*_. Accordingly, *G*^*a*^ is distributed as a binomial random variable with *K* trials and success probability *a*^*k*^. We denote this distribution as Binom(*a*^*k*^, *K*).

Our goal, as before, is to determine whether *a* ≥ *A*, which is equivalent to testing whether *a*^*k*^ ≥ *A*^*k*^. We can thus reduce the problem to one of testing whether *a*^*k*^ ≤ *A*^*k*^ based on a single sample |𝒮 |^*a*^ ∼ Binom(*a*^*k*^, *K*). This is a classical problem of hypothesis testing, a field famously studied by Neyman and Pearson [38]. Neyman and Pearson identified that to appropriately perform hypothesis testing, one must separate errors into two types: type I errors, or false negatives, in which the test incorrectly rejects a null hypothesis (*H*0) that is actually true; and type II errors (false positives), in which the test fails to reject a null hypothesis which is false. Accordingly, the established approach when constructing such tests is to choose a *significance level* (one minus the probability of error) for one of the two types, and then to endeavor to minimize the probability of the other.

We pursue such an approach here, choosing to specify the probability of false negative (the failure to report the presence of an organism in the sample). To do this, we define the *hypothesis boundary* as follows:

#### Definition 3 (Hypothesis Boundary)

*The* ***hypothesis boundary*** *for n k-mers with k-mer size k, ANI threshold A, and significance level α is denoted q*(*n, k, A, α*) *and is defined as the largest integer q satisfying* ℙ (*X* ≥ *q*) ≥ *α for a random variable X* ∼ *Binom*(*A*^*k*^, *n*).

In biological terms, given a set of *k*-mers *G, q*(|*G*|, *k, A, α*) is the greatest number such that the sketch of a randomly mutated copy of *G* with ANI exactly *A* will contain at least *q*(|*G*|, *k, A, α*) unmutated *k*-mers with probability at least *α*. Therefore, by definition, given significance level *α*, the test that rejects the null hypothesis whenever | 𝒮 |*< q*(|*G*|, *k, A, α*) and accepts it otherwise, is guaranteed to have false negative rate at most 1 − *α*; see Algorithm 1. Moreover, this holds true with no distributional assumptions on the ANI *a*^1^.

Biologically speaking, we claim that our reference genome is present whenever the sample shares at least *q*(|*G*|, *k, A, α*) *k*-mers with the reference genome; when fewer than *q*(|*G*|, *k, A, α*) shared *k*-mers are present in the sample, we conclude that too many *k*-mers have mutated for the sample genome to be related to the reference at the *A* ANI level. In the next section, we also take into consideration “coverage”, that is, when only a fraction of a genome is present in a given sample, modeling situations where insufficient sequencing depth resulted in reads covering only a portion of the genome.

#### Algorithm 1

YACHT_N1, *N* = 1

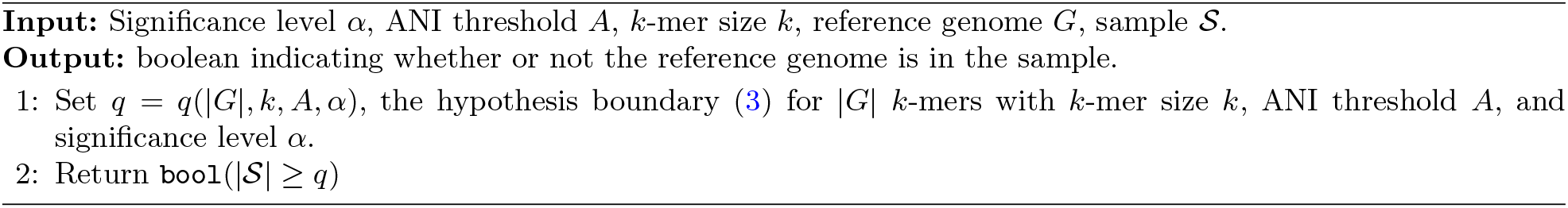

### 2.4 Accounting for Coverage

The model proposed so far assumes *k*-mers are absent from 𝒮 only when mutations occur that remove them from the sketch. This neglects the possibility that the number of reads in the sample may not be sufficiently large to record every *k*-mer for every genome at least once. Effectively, the algorithm proposed above assumes a coverage (proportion of *k*-mers in genome recorded in the sample) of 1, which cannot be guaranteed in practice.

One can see that accounting for coverage will require tradeoffs in terms of false negative and false positive rate: for instance, to guarantee a fixed rate of false negatives in the presence of arbitrarily small coverage would require accepting the presence of organisms sharing even a single *k*-mer with the sample. To avoid introducing distributional assumptions on the coverage parameter, we introduce a user-specified minimum coverage parameter *C* ∈ (0, 1] and calibrate the hypothesis boundary in Algorithm 1 to guarantee significance *α* for any organism in the sample that has at least a *C*-fraction of its exclusive *k*-mers present in the sample. The resulting algorithm, detailed in Algorithm 2, is the same as Algorithm 1 with |*G*| replaced by ⌊*C*|*G*|⌋ in the computation of the hypothesis boundary *q*, where ⌊·⌋ is the integer floor function.

#### Algorithm 2

YACHTC_N1, *N* = 1

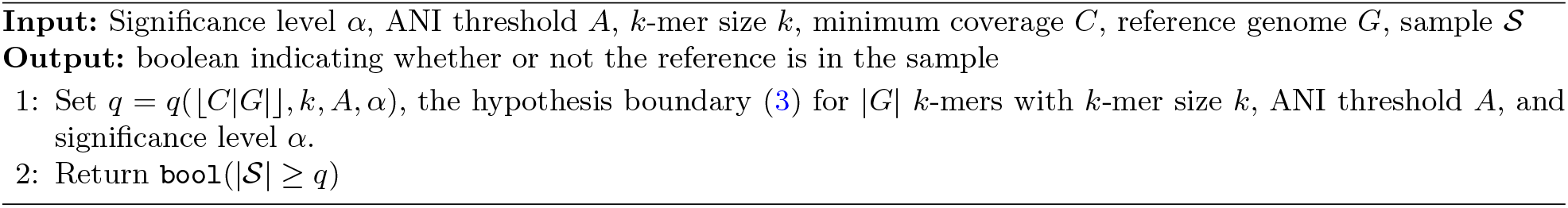

### 2.5 YACHT: General Case, *N* ≥ 1

In the general case, we cannot employ the same algorithm as in the *N* = 1 case due to the issue of shared *k*-mers between genomes. Unmodified application of Algorithm 2 would allow for the appearance of *k*-mers in the sample to support the presence of multiple organisms in the reference, leading to potential false positives.

However, due to the properties of metagenomic data, we are able to employ Algorithm 1 with only minor modifications. In practice, the sketches even of closely related genomes will still contain a significant proportion of *k*-mers exclusive to each genome (that is, *k*-mers which are not contained in any other genome sketch *G*_*i*_), as long as *k* is sufficiently large. Thus we can restrict application of Algorithm 2 only to those *k*-mers which are not shared by multiple genomes in the reference. Mathematically speaking, we employ the following definition:

#### Definition 4 (Exclusive *k*-mers)

*Given a collection of genomes* 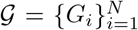, *the set* 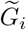 *of* ***exclusive*** *k****-mers*** *of genome i is defined as the set of k-mers in G*_*i*_ *that are not in G*_*j*_ *for any j* ≠ *i:*

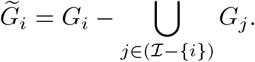

We then apply Algorithm 2 to 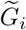 and the restricted sample 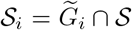. By only considering exclusive *k*-mers, the decision to include each genome has no effect on the inclusion of the others, allowing us to perform the hypothesis test independently and iteratively on each. By the assumption of independence on mutations between organisms, this has no effect on the accuracy of tests for individual organisms (aside from the effects of testing with fewer *k*-mers due to only using each organisms’ exclusive *k*-mers).

To maximize the number of exclusive *k*-mers in each genome, we first pre-filter our reference by eliminating any genomes that share no *k*-mers with the sample. These genomes will never be judged present, and so are removed to prevent any possible shared *k*-mers from interfering with genomes which are not filtered out. The result is the YACHT algorithm detailed in Algorithm 3. We note that algorithm 3 assumes the same minimum coverage for every organism, but there is no reason this parameter cannot be specified per organism.

### 2.6 Statistical Power

Since the probability of false negative error is generally fixed by the test design, the appropriate measure of the effectiveness of a hypothesis test is its *power* : the probability that, given the null hypothesis is false, the test actually rejects the null hypothesis. We note that, although the probability of false negative is fixed by the parameter *α*, the power of YACHTC N1 will vary between organisms in the reference depending on the number of exclusive *k*-mers each has, with organisms with more exclusive *k*-mers enjoying a lower probability of false positives.

#### Algorithm 3

YACHT

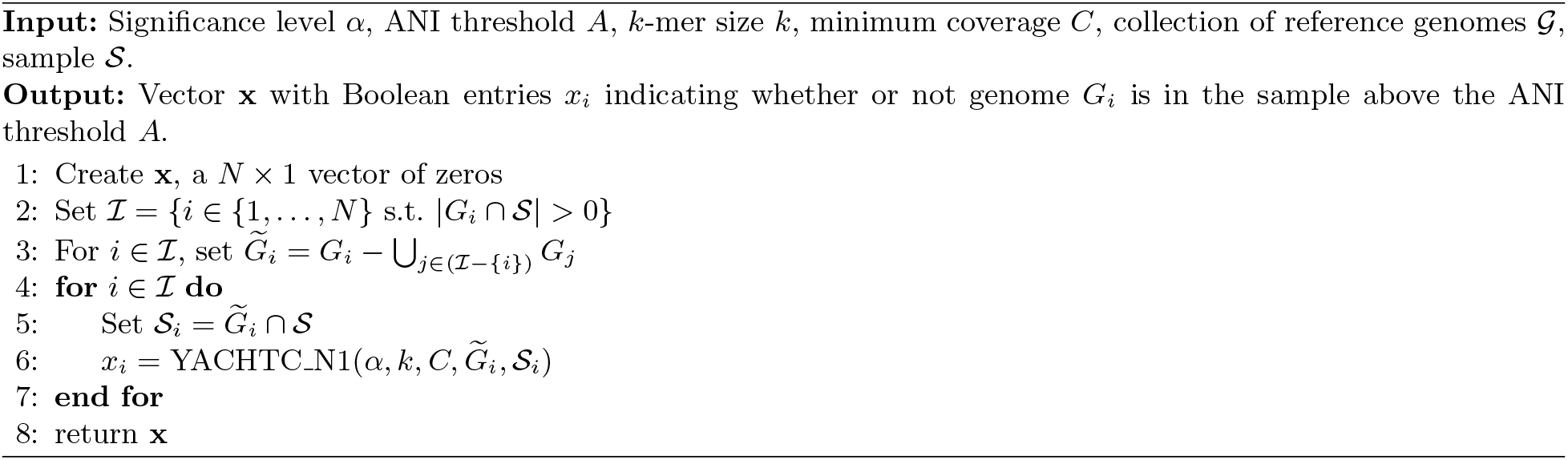

Computing YACHT’s power exactly would require a precise distribution for ANI’s, which we have deliberately avoided due to this distribution being highly dependent on the biological situation to which this algorithm is applied. As a distribution-free proxy, we can instead compute the *alternative significance* 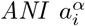:

#### Definition 5 (Alternative Significance ANI)

*Given a collection of genomes 𝒢, sample 𝒮, and significance level α, the* ***alternative significance ANI*** *for organism i, denoted* 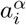, *is the unique solution to the equation*

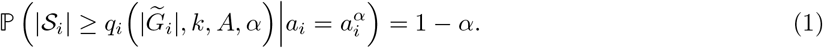

*In other words*, 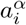 *is the ANI such that if* 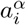 *were the true ANI of organism i, the probability of YACHT returning a false positive is exactly* 1 − *α*.

Equation 1 can easily be solved numerically due to the monotonicity of 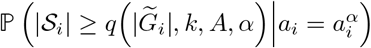 as a function of 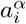. It is also the case that this monotonicity guarantees 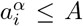, and it implies that any genome with an ANI less than 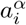 will be even less likely to appear as a false positive. The closer 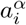 is to the ANI threshold *A*, the narrower the range in which YACHT is likely to make a false positive judgement and so the greater the statistical power.

### 2.7 Practical Considerations

Algorithm 3 can be run for any set of reference genomes 𝒢. In practice, however, good performance will only be achieved as long as each genome has an appreciable number of exclusive *k*-mers, which requires that no two reference genomes share essentially all of their *k*-mers. Accordingly, we employ the following pre-processing algorithm to ensure that no two organisms are more closely related than the species ANI threshold *A*. By a greedy algorithm, we find a *A-distinct subcollection* 𝒢^*A*^ ⊆ 𝒢 defined as follows:

#### Definition 6 (*A*-distinct subcollection)

*Given k-mer size k, ANI threshold A, and collection of reference genomes* G, *an A****-distinct subcollection*** *of* 𝒢 *is a collection* 𝒢^*A*^ ⊆ 𝒢 *such that for every* 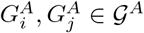 *with i* ≠ *j:*

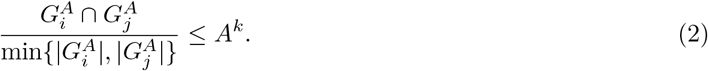

In biological terms, whenever two reference organisms are related above the ANI threshold, we choose one to represent that “class” of related organisms. We also design our algorithm to reject the smaller of any two related genomes, as the larger will typically enjoy greater statistical power with YACHT. The exact algorithm is detailed in Algorithm 4.

The effects of excluding such a preprocessing step are reflected in figure 8, which for synthetic data shows a dramatic increase in false positive rate when organisms in the reference are more closely related than the ANI threshold.

#### Algorithm 4

YACHT Preprocessing

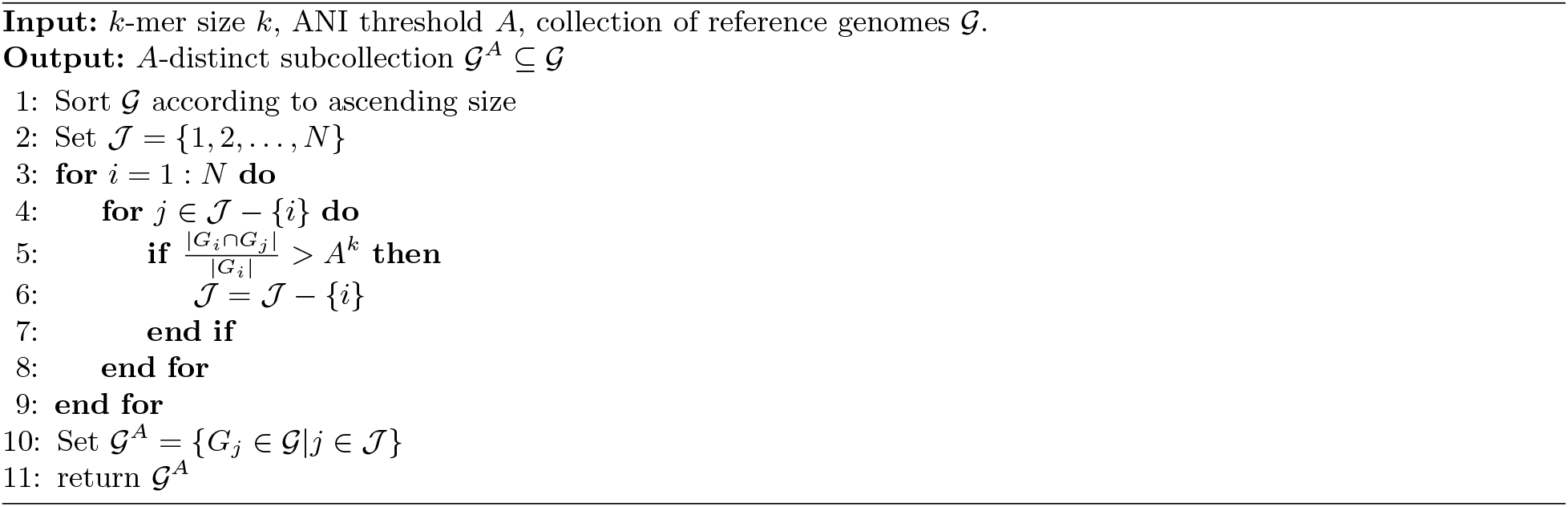

## 3 Theory

In this section, we provide theoretical bounds on the hypothesis boundary *q*(*n, k, A, α*) as well as the alternative significance ANI *a*^*α*^. For simplicity of notation we restrict consideration to the single-genome case, but our results apply equally to the *N >* 1 case under the substitution 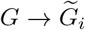 and 𝒮 → 𝒮_*i*_. To further simplify notation, in this section we will use the following shorthand: *n* = |*G*|, *q* = *q*(*n, k, A, α*), and *μ* = *nA*^*k*^. In both of our theorems, we make use of Chernoff bounds on sums of Bernoulli random variables. Variants of these are well-known in probability literature; a more precise version, of which the following theorem is a corollary, is proven in [39, Theorem 4.4].

### Theorem 1

**(Chernoff Inequality)***Let X*_1_, …, *X*_*N*_ *be independent random variables taking values in* {0, 1} *with* ℙ (*X*_*i*_ = 1) = *p*_*i*_. *Then setting* 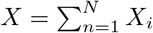 *with* 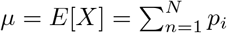 *and δ >* 0, *we have:*

1. *If δ* ∈ (0,1], *then* ℙ(*X* ≥ (1 + *’*) µ ≤ exp (−*µδ*^2^/3)
2. *If δ* ∈ (0,1], *then* ℙ(*X* ≥ (1 − *’*) µ ≤ exp (−*µδ*^2^/3)
3. *If δ* < 1,*then* (*X* ≥ (1 + *’*) µ ≤ exp (−*µδ*^2^/3)

We begin with the following theorem, which bounds the distance between *q* and the mean number of unmutated *k*-mers *μ*:

### Theorem 2

*Fix α* ∈ (0, 1). *Let X be a binomial random variable with probability of success p, n trials, and mean μ* = *nA*^*k*^. *Let q be the largest integer such that* ℙ (*X* ≥ *q*) ≥ *α. Then there exists a positive constant* 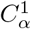 *depending only on α such that:*

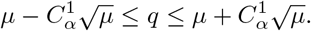

*Proof*. We can equivalently define *q* as the largest integer such that ℙ (*X < q*) ≤ 1 − *α*. By Theorem 1.2,

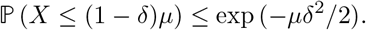

Setting exp(−*μδ*^2^*/*3) = 1 − *α*, we can compute that ℙ (*X* ≤ (1 − *δ*)*μ*) ≤ 1 − *α* so long as

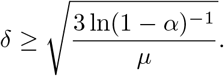

We conclude that

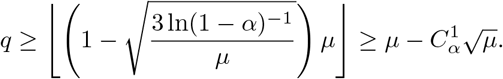

Similar logic for the upper tail gives 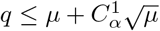, possibly changing the constant 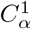.

As a corollary, we can immediately infer that as *n* becomes large, *q* tends to be close to the mean *μ*:

### Corollary 1

*μ/q* → 1 *as n* → ∞.

We now prove a bound relating the alternative significance ANI *a*^*α*^ to the ANI threshold *A*:

### Theorem 3

*Fix α* ∈ (0.5, 1) *and let q be the largest integer such that* ℙ (*X* ≥ *q*) ≥ *α for X* ∼ *Binom*(*A*^*k*^, *n*). *Let a*^*α*^ *be the number in* (0, *A*) *such that for Y ~ Binom*((*a*^*α*^)^*k*^, *n*), ℙ (*Y* ≥ *q*) = 1 − *α. Then there exists a positive constant* 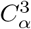 *depending only on α such that:*

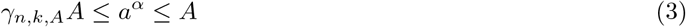

*where*

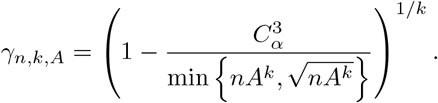

*In particular, as n* → ∞ *with k fixed, a*^*α*^ → *A*.

In words, this means that the statistical power of the test approaches 1 as the number of exclusive *k*-mers increases.

*Proof*. We have already concluded that *a*^*α*^ ≤ *A*, and so proving the result requires proving only the lower bound. To accomplish this, we set *μ*_*α*_ = *n*(*a*^*α*^)^*k*^ and employ Theorem 1.1-3, which yields for *δ* ∈ (0, 1]

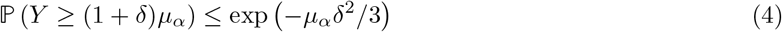

and when *δ* > 1,

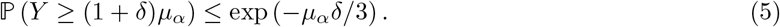

We divide into the cases *δ* ≤ 1 and *δ* > 1, which corresponds to the cases (*a*^*α*^)^*k*^ ≤ 2*A*^*k*^ and (*a*^*α*^)^*k*^ > 2*A*^*k*^. We begin with the case (*a*^*α*^)^*k*^ ≤ 2*A*^*k*^. By the change of variables (1 + *δ*)*μ*_*α*_ → *q*, Theorem 1.2 gives:

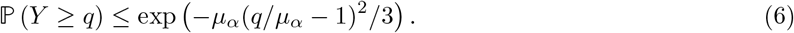

Next, we claim that for some positive 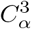 depending only on *α*,

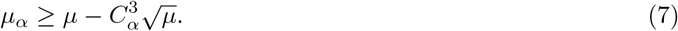

We consider two cases. First, suppose that *μ*_*α*_ ≥ *q*. Then by theorem 2,

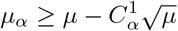

so the claim holds with 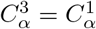.

Next, we consider the case *μ*_*α*_ *< q*. We set the right-hand side of (6) to 1 − *α*, yielding

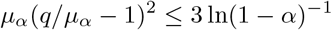

which as an equation has two solutions for *μ*_*α*_:

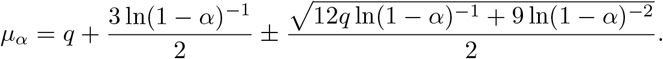

Since *μ*_*α*_ < *q* by assumption, we know that the lower bound for *μ*_*α*_ must be the smaller of these two solutions, as the other will be greater than *q*. Accordingly,

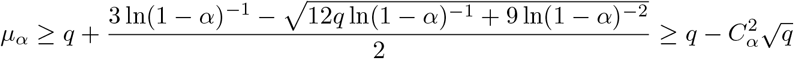

where 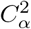 is a constant depending only on *α*; thus claim (7) holds for all *μ*_*α*_.

Substituting the result of theorem 2 into (7), we have

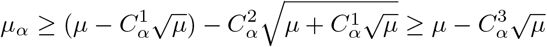

for new constant 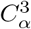 depending only on *α*. Substituting *μ*_*α*_ = *n*(*a*^*α*^)^*k*^ and *μ* = *nA*^*k*^, we have

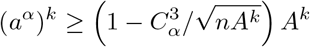

and therefore

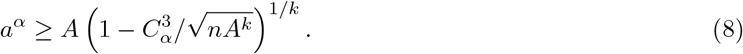

as long as (*a*^*α*^)^*k*^ ≤ 2*A*^*k*^.

Following essentially the same steps in the case (*a*^*α*^)^*k*^ > 2*A*^*k*^ and applying Theorem 6.1, we find that in this case

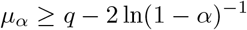

and thus that

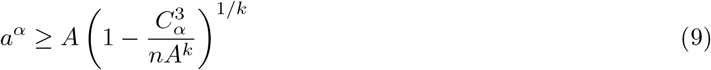

for possibly changed 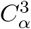. Taking the larger of the bounds from equations (8) and (9) completes the proof.

Theorem 3 also suggests the somewhat surprising result that increasing *k* does not monotonically improve accuracy (in the sense of bringing *a*^*α*^ closer to *A*). The *γ*_*n,k,A*_ term in line (3) has a unique global and local maximum for *k* ≥ 1, suggesting there is an optimal *k* that maximizes the power of `YACHT` for a specific number of unique *k*-mers *n*. This phenomenon corresponds to the fact that for large *k*, nearly all of a genome’s *k*-mers will be mutated at the ANI threshold *A*, so the hypothesis boundary *q* must be set very close to zero. When this happens, only small deviations are necessary for more highly mutated genomes to cross the inclusion threshold. This behavior is reflected in the following section in Figure 3.

The critical point could be estimated theoretically using the estimates of 3, but as these are only approximations it will be more practical to simply test a range of *k*-mer values with the exact binomial CDF. It should also be noted that this analysis assumes *n* remains unchanged as *k* increases, when in practice the number of *k*-mers unique to each organism can be expected to increase with *k* as overlaps become less probable, which would partially mitigate the downside of increased *k* in real-world applications.

## 4 Synthetic experiments

In this section, we describe two sets of experiments: one on synthetic data to aid in understanding the impact on performance of various model parameters, and one on real-world data to test the correspondence of the theory with reality.

### 4.1 Synthetic Data

In this section, we show results of the `YACHT` method using simulated data. The simulation parameters are detailed in Table 1. When not specified as part of the experiment, the default parameters are used. We note that our synthetic data always has in-sample coverage of 1; see section 5.2 for experiments verifying performance when true coverage is less than one. All simulations were run for 100 iterations for each set of parameters.

**Table 1:** Synthetic data simulation parameters.

| Parameter Name | Symbol | Description | Default |
| --- | --- | --- | --- |
| --ksize | $k$ | $k$-mer size | 31 |
| --num_kmers | $n$ | Number of $k$-mers per genome | 1000 |
| --num_genomes | $N$ | Number of genomes in reference | 1000 |
| --s_known | $s_k$ | Number of true known organisms in sample | 100 |
| --s_unknown | $s_u$ | Number of true unknown organisms in sample | 100 |
| --ani_thresh | $A$ | Mutation rate cutoff for known/unknown | 0.95 |
| --relation_thresh | $r$ | Maximum ANI between reference genomes | 0.95 |
| --significance | $\alpha$ | Significance level for testing | 0.99 |
| --min_coverage | $C$ | Minimum coverage for significance level | 1 |
| --ani_range | $[m, M]$ | Range of possible ANI's | $[0.9, 1]$ |

We note that we have set the default minimum ANI to 0.9, which is much lower than the largest mutation rates seen in practice. As will be shown, YACHT only experiences false positives relatively close to the ANI cutoff *A*, so a low upper bound was chosen to ensure a nontrivial number of false positives in the resulting simulations. The choice of 0.9 exactly allows for symmetric distribution of mutation rates around the ANI threshold *A* = 0.95. As a result of this choice, the observed false positive rates are inflated relative to what might be expected in practice, where the lowest true ANI’s will be closer to 0.7. To aid in interpretation of our numerical results, roughly speaking, in this setting a 0.2 false positive rate corresponds to accepting false positives with true ANI as low as 0.94, while a 0.5 false positive rate would correspond to accepting false positives with true ANI as low as 0.925.

#### Simulated Reference Model

We employ the following model for the reference genomes 𝒢. The total number of *k*-mers across all genomes is set at *K* = *nN* (1 − *r*^*k*^) (rounded to the nearest integer). The first genome *G*_1_ consists of *n* randomly selected *k*-mers, while subsequent genomes *G*_*i*_ consist of ⌊*nr*^*k*^⌋ *k*-mers chosen uniformly from *G*_*i−*1_ and *n* − ⌊ *nr*^*k*^ ⌋ chosen uniformly from its complement. This model guarantees each genome in is highly related to at least one other genome in the reference with an ANI of *r*.

All simulations are run under this model except for runtime/memory simulations, which use a similar but deterministic model for the reference which can be generated more quickly. This does not affect the recorded recovery runtimes, which do not take into account the time spent setting up the synthetic data.

#### Simulated Sample Model

The sample 𝒮 is constructed according to the random sample model 2 with randomized mutation rates **r** set as follows. Out of the *N* genomes, a uniformly random set of *s*_*k*_ genomes is labeled known and each chosen genome is assigned an independent and uniformly random ANI in the interval [*A, M*]. Likewise, *s*_*u*_ genomes are randomly chosen to be unknown and each assigned an independent and uniformly random ANI in in [*m, A*]. The remainder of the genomes in 𝒢 are omitted from the sample (or, equivalently, assigned an ANI of 0).

### 4.2 Significance level reflects true false negative rate

Figure 1 shows the true false negative rate out of 200 known organisms when mutation rate is fixed at exactly 0.05. We see that false negative rate is always slightly less than 1 minus the specified significance level. As the binomial distribution is discrete, the hypothesis boundary cannot be chosen exactly to result in a false negative rate of 1 − *α*, and so is always an underestimate. Comparing the results for *n* = 1000 and *n* = 3000, we see that increasing the number of *k*-mers per genome reduces this difference.

**Fig. 1:**
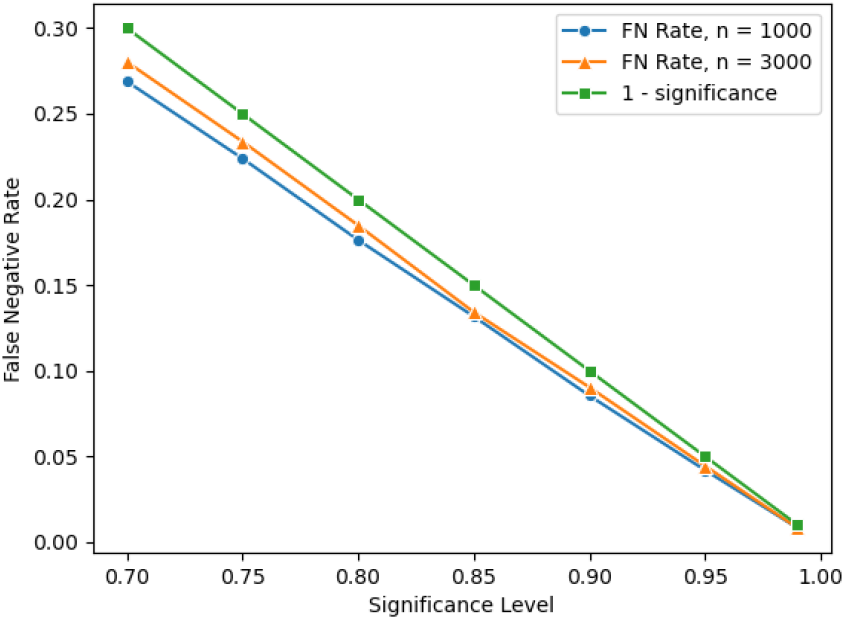
False negative recovery rate vs. significance level. The vertical axis gives the number of false negatives divided by the total number of known organisms over 100 iterations. The horizontal axis gives the value of the user-specified significance level *α*.

### 4.3 Prediction Rate vs. ANI

In Figure 2, we show the rate of positive results (organisms marked present in the sample) versus the underlying mutation rate. For this experiment, 200 organisms were in the sample with true mutation rates *exactly* equal to a specified ANI which was varied from 0.92 to 0.97. We see that for ANI substantially below the ANI threshold, the positive rate is 0, before quickly increasing to 1 above the threshold. We also observe that the transition from 0 to 1 occurs more quickly for *n* = 2000 than *n* = 1000, reflecting the increase in statistical power that accompanies a greater number of *k*-mers per organism. The blue and orange vertical lines show the alternative significance ANI (*a*^*α*^) for 200 and 400 exclusive *k*-mers, respectively, showing that a greater number of *k*-mers corresponds to *a*^*α*^ closer to the ANI threshold.

**Fig. 2:**
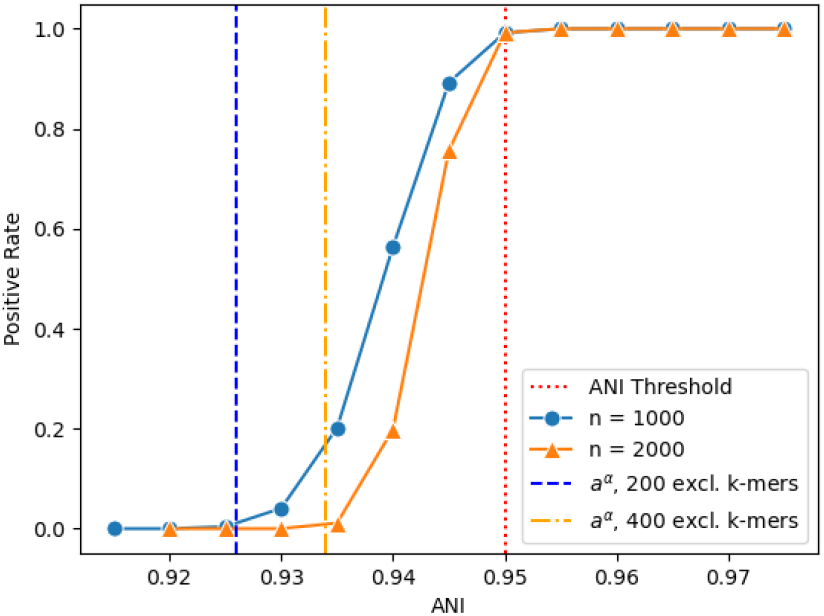
Positive rate vs. ANI. The vertical/*y*-axis gives the number of organisms marked present divided by the total number of organisms in sample over 100 iterations. The horizontal/*x*-axis gives the true ANI rate of each genome in the sample to a genome in the reference. The left-most and middle vertical dashed lines give the alternative significance ANI using 200 and 400 exclusive *k*-mers respectively; the right-most vertical dashed line gives the ANI threshold.

### 4.4 Performance in *k* reflects theoretical predictions

Figure 3 shows performance as *k*-mer size is increased. As an estimate for *a*^*α*^ for each simulated reference and sample, the minimum ANI among false positive results was tracked (with the ANI threshold *A* substituted in this average in the case of no false positives). Figure 3b shows the average of these minimum false positive ANI’s across 100 simulations for varying *k*-mer sizes *k*. As predicted by the *γ*_*n,k,A*_ term from Theorem 3, the minimum false positive ANI initially rises to a local maximum in *k* before decreasing once *k* gets too large. A similar pattern can be observed in the false positive rate in figure 3a.

**Fig. 3:**
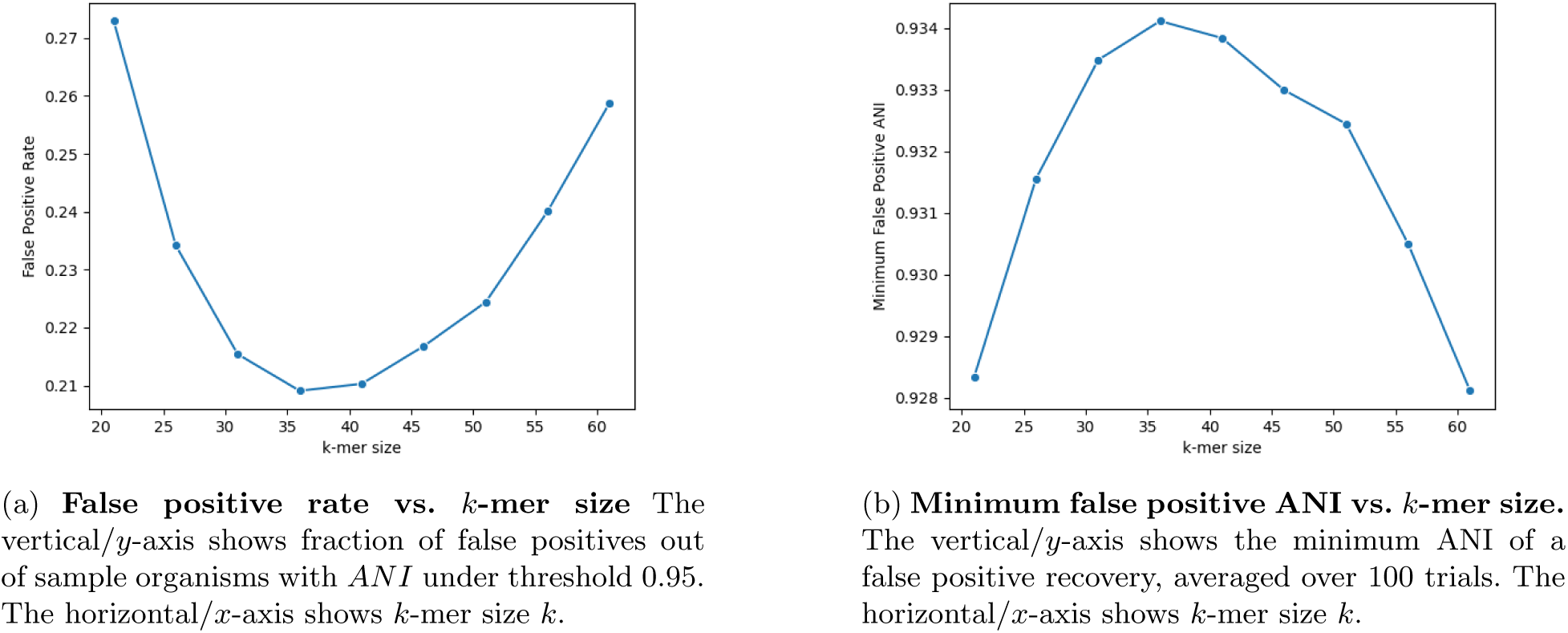
False positive performance vs. *k***-mer size**. Figure 3a (left) shows the effect of *k*-mer size on false positive rate, while Figure 3b (right) shows its effect on minimum false positive ANI.

### 4.5 Runtime and space usage

In Figures 4-5, we show how runtime and space usage vary with the two main parameters which affect it, number of *k*-mers per reference genome *n* and number of reference genomes *N*. Growth in runtime is at-most linear in both *n* and *N*, though the growth of runtime in *n* is somewhat inconsistent over small changes. By contrast, memory usage grows nearly exactly linearly in both variables.

**Fig. 4:**
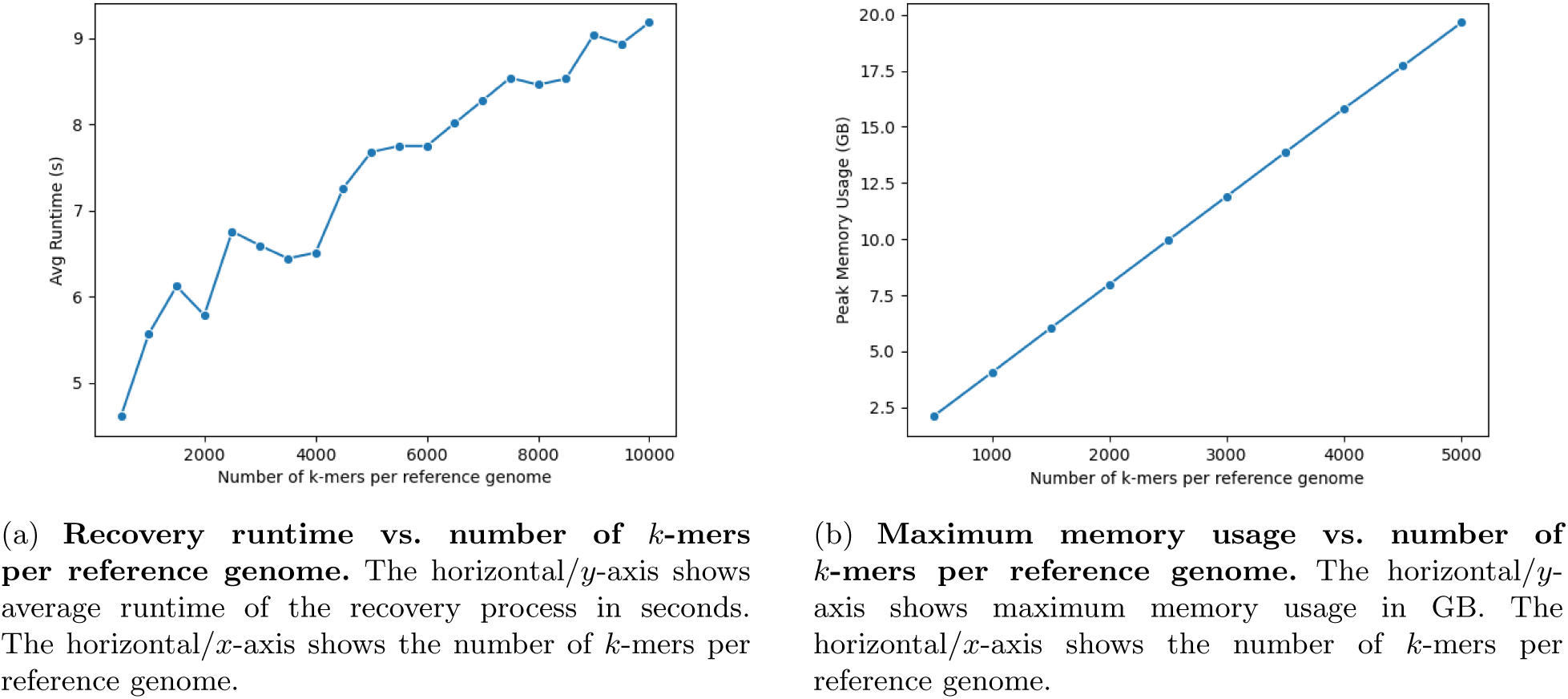
Runtime and memory usage vs. number of *k*-mers per reference genome. On the left, figure 4a shows the effect of number of *k*-mers per reference genome on recovery runtime while figure 4b on the right shows its effect on maximum memory usage.

**Fig. 5:**
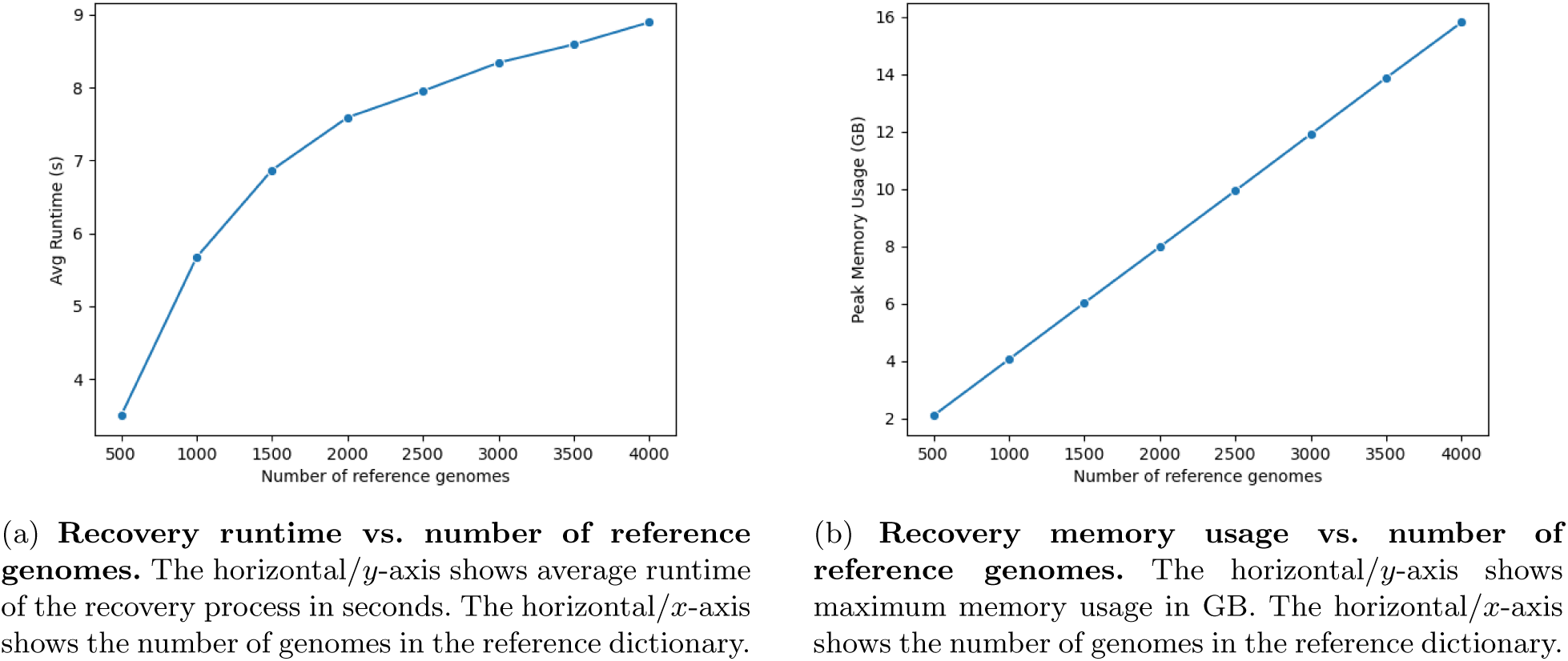
Runtime and memory usage vs. number of reference genomes. On the left, figure 5a shows the effect of number of reference genomes on recovery runtime while figure 5b on the right shows its effect on maximum memory usage.

### 4.6 Performance with Varying Parameters

In this section, we show how false positive and false negative rate are affected by changing various simulation parameters.

*N* : *Number of genomes in reference (figure omitted)*. In experiments with synthetic data, changing the number of genomes in the reference had essentially no effect on false positive or false negative rate. This is expected due to the use of exclusive *k*-mers implying each genome tested is independent from all others.

*c: Minimum coverage for significance level*. In Figure 7, we show how false positive rate changes with minimum coverage parameter *C*. As expected—given that true coverage remains unchanged at 1—a lower minimum coverage results in higher false positive rate.

**Fig. 6:**
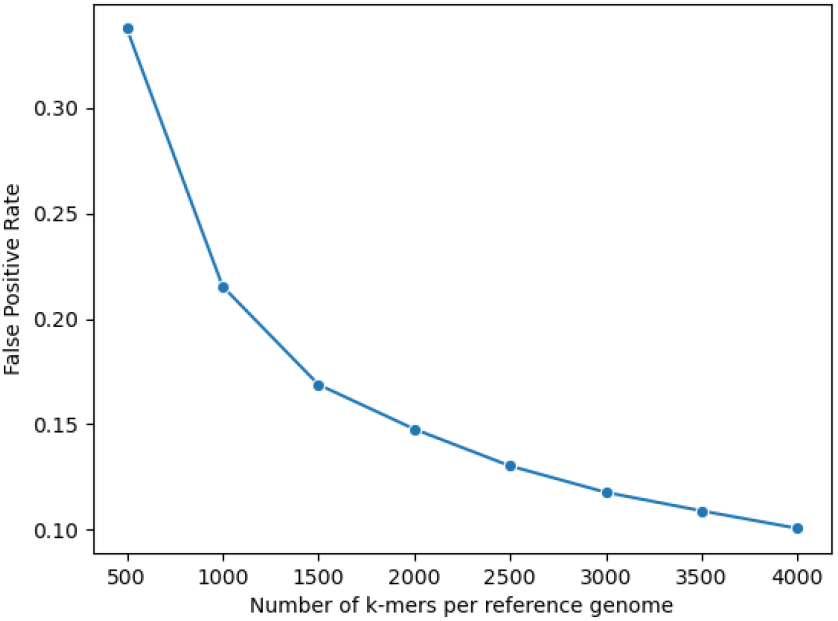
False positive rate vs. number of *k*-mers per reference genome. The vertical/*y*-axis shows fraction of false positives out of sample organisms with ANI under threshold 0.95. The horizontal/*x*-axis shows number of *k*-mers per genome in the reference dictionary.

**Fig. 7:**
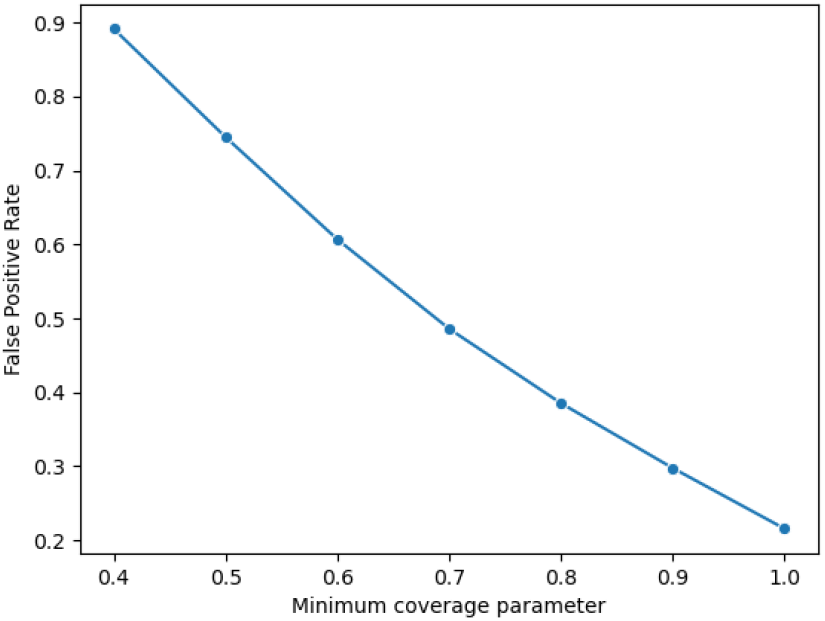
False positive rate vs. `--min coverage` **parameter** The vertical/*y*-axis shows fraction of false positives out of sample organisms with ANI under threshold 0.95. The horizontal/*x*-axis shows user-specified `--min coverage` parameter.

*r: Maximum ANI between reference genomes*. Figure 8 shows that as the ANI between reference genomes increases, so does false positive rate. An especially dramatic spike occurs when ANI exceeds the ANI threshold *A*, as this drastically reduces the number of *k*-mers unique to each genome in the reference.

**Fig. 8:**
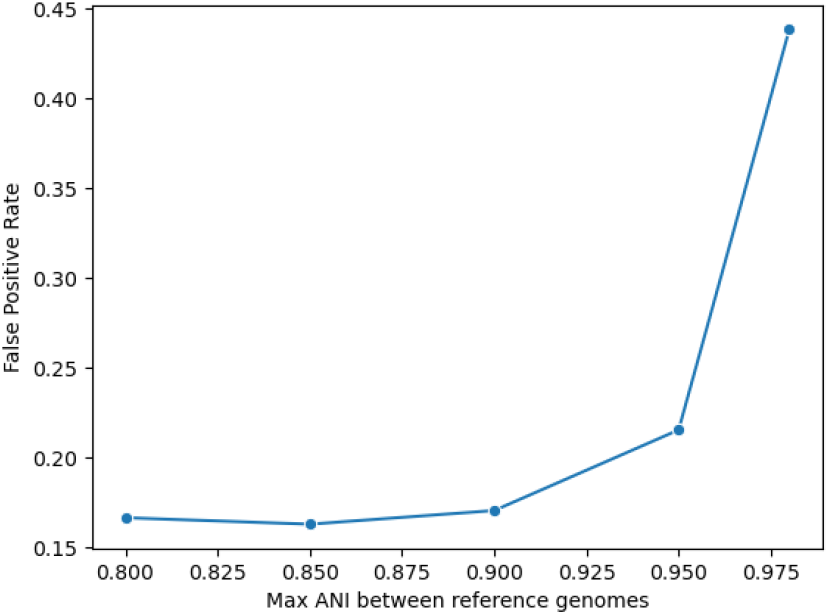
False positive rate vs. max ANI between reference genomes. The vertical/*y*-axis shows fraction of false positives out of sample organisms with ANI under threshold 0.95. The horizontal/*x*-axis shows the maximum ANI between reference genomes.

*α: Significance level*. Figure 9 demonstrates the tradeoff between false positive and false negative rate that is controlled by *α*. When *α* = 0.5, false positive and false negative rates are nearly equal; as *α* grows to 0.99, false negative rate declines nearly to zero while false positive rate becomes high.

**Fig. 9:**
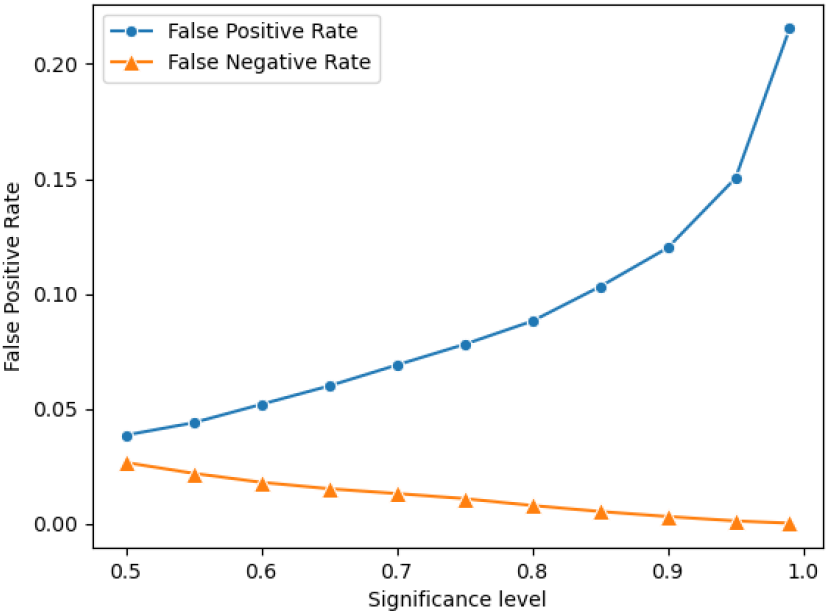
False positive/negative rates vs. significance level. The vertical/*y*-axis shows fraction of false positives and false negatives out of sample organisms with ANI under (respectively, above) threshold 0.95. The horizontal/*x*-axis shows the user-specified significance level *α*.

## 5 Real data experiments

For an end user biologist, the two parameters of main interest are the parameters `--min coverage`, denoted *C* (i.e. what percentage of a genome must be present for it to be detected) and `--ani thresh`, denoted *A* (i.e. how divergent a sample organism must be from a reference organism to be considered a “different organism”). As such, we consider each in turn for a series of “spike-in” experiments. In each experiment, (portions of) real genomes are mixed into a real metagenomic sample and `YACHT` is run in order to assess if this spiked-in genome can be detected as present. Before detailing the experiments, we describe the associated data.

### 5.1 Real data

#### Reference data

To form the **reference database** of genomes 𝒢, we collected 1,044 bacterial genomes randomly selected from NCBI RefSeq [40]. We utilized Sourmash [23] to form FracMinHash sketches of each of these genome using a *k*-mer size of 31 and a scale factor of *s* = 1*/*1, 000. In all subsequent experiments, the preprocessing from Algorithm 4 was run with the ANI threshold *A*.

#### Real metagenome

We selected a random, publicly available human gut metagenome from the ENA [41] which we call the **real metagenome**. Specifically, from the project PRJEB9576 we selected the Illumina HiSeq 2000 sample with accession number SAMEA3449194 and took the forward reads only, for a total of 27M reads. We utilized Sourmash’s `gather` metagenome profiler to predict which genomes from our reference database are present in this metagenome. Importantly, we set the parameter `--coverage-bp` to zero (i.e. maximum sensitivity) in order to detect the most reference genomes present in this metagenome. A total of 268 reference database genomes were detected as being “present” in this metagenome and we refer to the remaining 776 reference database genomes as **absent reference genomes** as they are, with high confidence, not present in the metagenome.

#### Known unknowns

In order to test the `--ani thresh` parameter, we need to collect genomes that are not contained in the reference data, but are at a known average nucleotide identity (ANI) to genomes that are in the reference. To that end, we downloaded the GTDB [37,36] R07-RS207 genomic representatives ^2^ database which contains 65,703 genomes. After forming FracMinHash sketches, we utilized the approach of [22] via Sourmash to compute the ANI of each of these 65K genomes to the absent reference genomes described in the previous paragraph, retaining those with an ANI above 0.7 to any of the absent reference genomes. We then selected from these those genomes that were absent from the real metagenome in the exact same way as in the previous paragraph. This resulted in a total of 28,595 **known unknowns**: genomes that are not in the sample nor in the reference, but are at a known ANI to genomes in the reference. These will be the spike-in genomes used in Section 5.3.

### 5.2 Coverage parameter accurately respects real coverage

Herein, we aim to test if the parameter `--min coverage` allows `YACHT` to successfully recover a genome from a sample (respectively, correctly report it is absent from the sample) if the coverage is sufficiently high (respectively, if the coverage is too low). In order to quantify this, individually for each of the above described “absent reference genomes”, we selected error-free, mutation-free reads from it until a total coverage *c* was attained. We then inserted these reads into the metagenome and ran `YACHT` using a `--min coverage` value of *C*. The spike-in genome was then labeled “detected” or “not detected” based on the output of Algorithm 3. We then varied the values of *c* and *C* by powers of two and repeated this experiment 100 times, varying the absent reference genomes and then averaging results. Figure 10 displays the results as a heatmap. As Figure 10 demonstrates, our approach successfully recovers a genome from a metagenome if its coverage in the metagenome is at or above the set `--min coverage` value with very high probability. The correspondence of the actual coverage of the genome in the metagenome and the `--min coverage` parameter is not perfect though, as some percentage of spike-in genomes were detected but had coverage in the sample lower than the set parameter. However, the relationship should be viewed as a lower bound of detection: setting `--min coverage` to some value *C* all but guarantees the ability to detect a genome with actual coverage at least *C* in the metagenome.

**Fig. 10:**
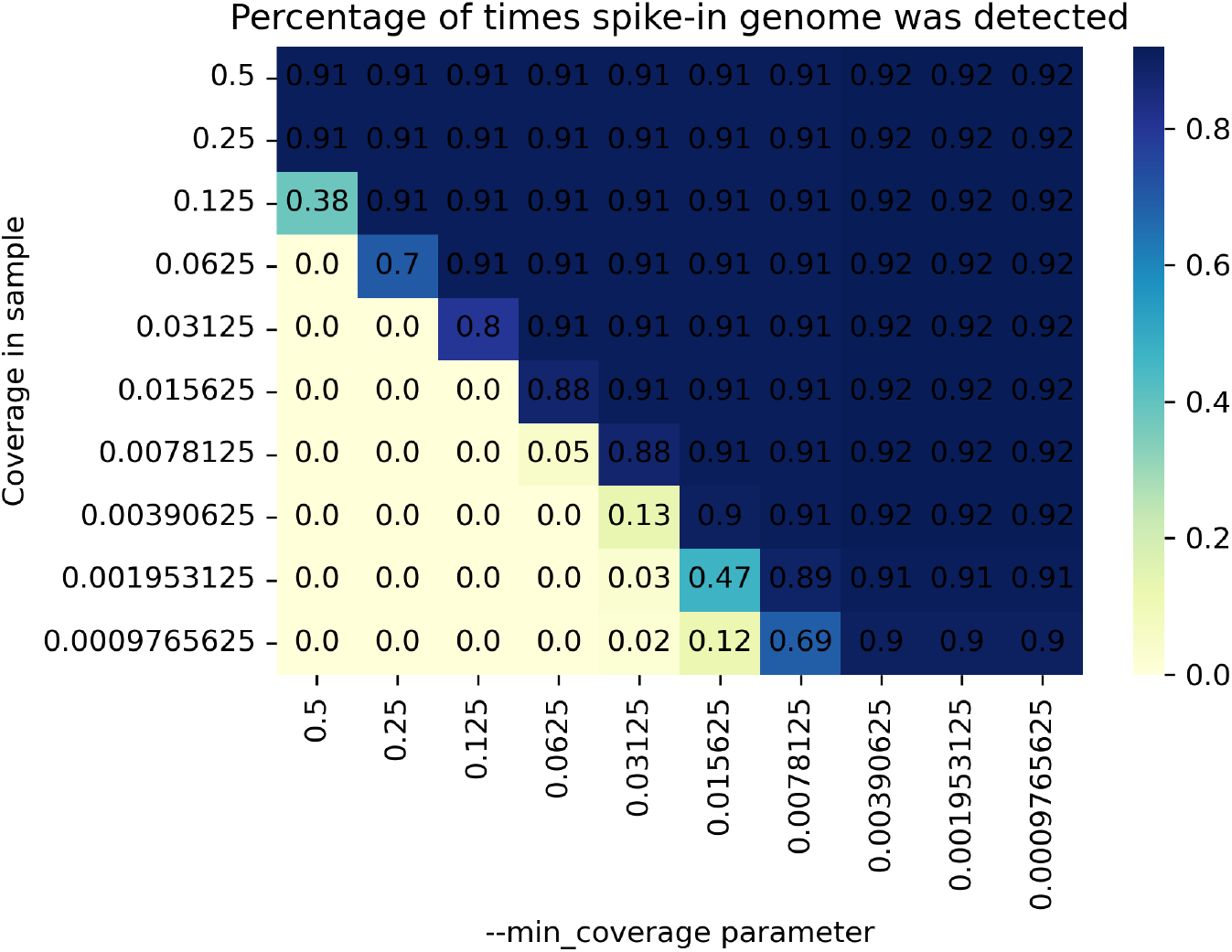
Heatmap of percentage of times the spike-in genome was recovered from the metagenome. The vertical/*y*-axis gives the coverage of the spike-in genome in the metagenome, called *c* in the main text. The horizontal/*x*-axis gives the value of the parameter `--min coverage`, called *C* in the main text. Both axes vary by powers of two.

### 5.3 Mutation rate spike in experiments

To assess the impact of the `--ani thresh` parameter, we utilize the “known unknown” genomes and spike them in the real metagenome similar to the previous section. First, we fixed the coverage of each spiked-in genome to *c* = 1 and set the `--min coverage` to *C* = 1. We then created 28,595 metagenomes, one for each “known unknown” spiked into the real metagenome, and ran `YACHT` on each metagenome. For each of these, we recorded if a similar organism was detected or not: that is, if Algorithm 3 detected an genome *G*_*i*_ as present in the sample, where *G*_*i*_ is the reference database genome with maximal ANI to the spiked-in “known unknown” genome. Of note, the actual relative abundance of the spike-in genomes was at most 0.35% on the basis of reads in the metagenome and from the spike-in genome.

Figure 11 shows the results of this experiment. In more detail, this figure shows that in 97.06% of these experiments, a similar organism was correctly not detected (a true negative) due to the ANI of the spiked organism to the reference being lower than the `--ani thresh` value, as seen by the blue dots on the bottom, left of the vertical red dashed line. A total of 99.80% of these experiments, `YACHT` correctly detected the reference organism similar to the spiked organism (a true positive), the ANI of this pair being above the `--ani thresh` parameter, as seen by the blue dots in the top, right of the vertical red dashed line. The false positive cases where the spiked organism had an ANI to the reference genome *G*_*i*_ below the ANI threshold, yet `YACHT` still detected *G*_*i*_ can be seen in the top to the left of the vertical red dashed line. We hypothesize that these are due to *k*-mers in the sample that match with those in *G*_*i*_ though originating from organisms different from the spike genome. The slightly increased false negative rate of 0.029 over the significance level, which was here set to 0.01 is likely due to violations of our assumptions in Section 2.2 due to real-world mutations not being independent nor one-way.

**Fig. 11:**
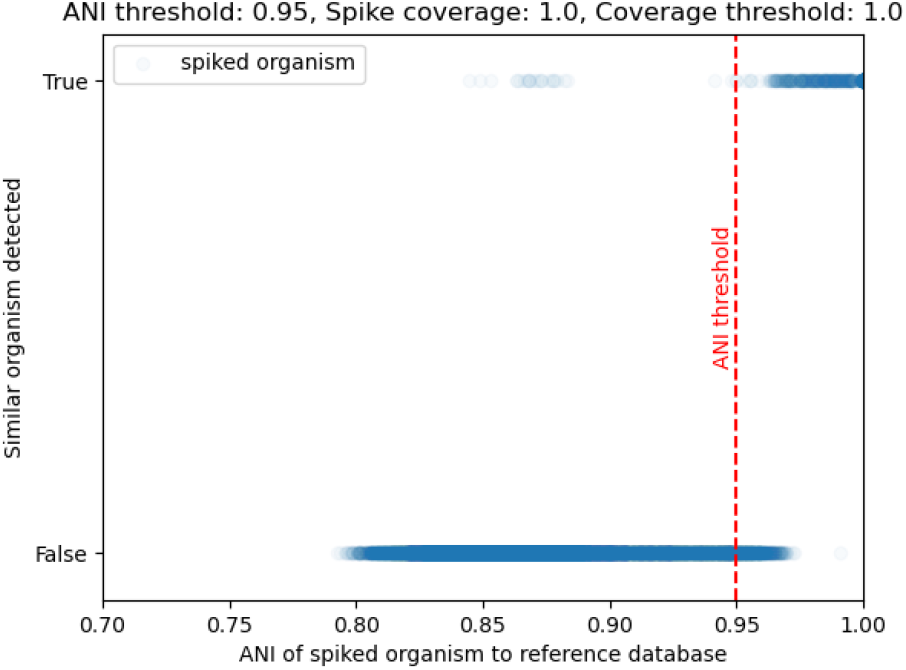
Scatter plot of 28,595 metagenomes, each with a single spiked-in genome absent from the reference database, but with a known maximal average nucleotide identity (ANI) to a reference database genome *G*_*i*_. This ANI value is recorded on the *x*-axis. The binary *y*-axis records if `YACHT` reported *G*_*i*_ as present (True) or absent (False) and a pale blue dot is recorded accordingly. The red vertical line depicts the setting of the parameter `--ani thresh`, here set to an ANI value of *A* = 0.95.

Next, we varied both the `--min coverage` parameter *C* and the coverage of the spiked-in genome in the metagenome *c*. The experiment was performed in the same fashion as before in Section 5.3, but here we record only the false positives and false negatives for ease of visualization. Figure 12 shows the result of these experiments when *c* and *C* both vary over 4 orders of magnitude, from 1.0 to 0.001. Of note, the lowest relative abundance of a spike-in genome was .00035% in the lowest *c* case. As these figures demonstrate, with decreasing coverage threshold *c* and/or decreasing `--ani thresh`, both false positives and false negatives increase. This is expected due to a coverage threshold of *c* = 0.001 corresponds to `YACHT` needing to see approximately 10^*−*6^ of a genome’s unique *k*-mers to predict it as “present” (due to the scale factor of *s* = 1*/*1000 and *c* = 0.001). For many organisms, this can be as low as a single *k*-mer or even no *k*-mers if that organism’s genome is small enough. As such, controlling the false positive rate can be had by either increasing the coverage threshold, or else increasing the scale factor.

**Fig. 12:**
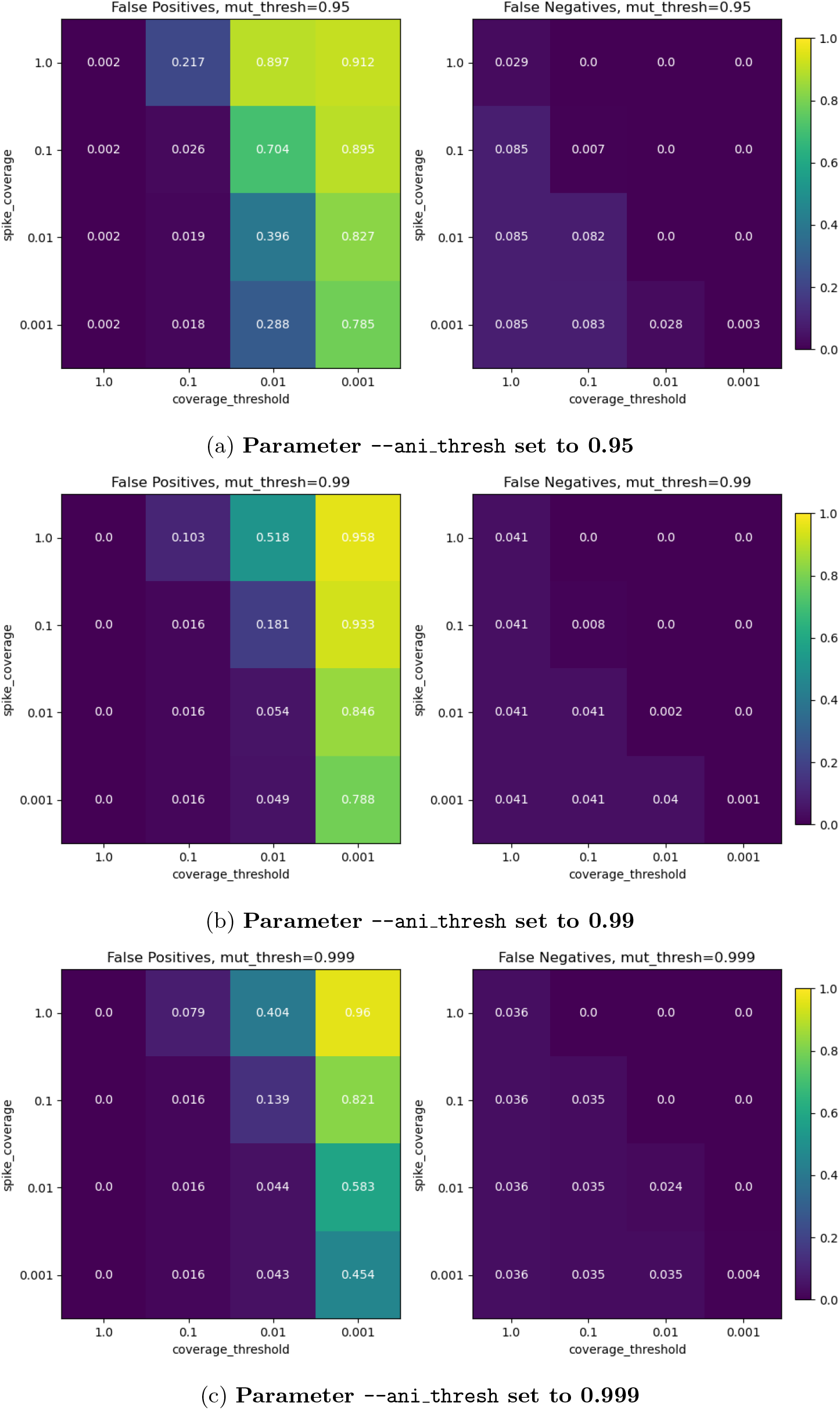
False positive and negative rates for spike in experiments. The heatmaps on the left show the False positive rate and the ones on the right show the false negative rate when the `--ani thresh` parameter was set to (a) 0.95, (b) 0.99 and (c) 0.999. In each heat map, the horizontal axis varies the coverage threshold parameter *C* and the vertical axis varies the coverage *c* of the spiked in genome in the metagenome.

### 5.4 Comparison with Sourmash

The tool Sourmash [23] possesses a sub-routine called `prefetch` which performs organism presence/absence detection via FracMinHash calculated containment indices. Sourmash’s subsequent sub-routine `gather` then uses a minimum set cover approach to reduce the number of false positives. In this section, we compare the performance of this presence/absence approach versus `YACHT`.

Due to needing a ground truth in order to assess performance, we used a mix of real and simulated data. For the sample metagenomes, we randomly selected 200 genomes from the **reference database** described above and used `BBMap` [42] to generate a metagenome from them consisting of 10 million noisy reads. To mimic the situation where the metagenome contains novel genomes not present in the reference database, we then removed half (100) of these genomes from the reference database before running Sourmash `prefetch` and `gather` (with a `--threshold-bp` value of 100 and all other parameters at their defaults) and `YACHT` (with a `--min coverage` value of 1 and all other parameters at their defaults). This procedure was then repeated a total of 30 times each for two different distributions of genome relative abundance: uniform and exponential distributions. The resulting false positive and negative rates are depicted in Figure 13. As can be seen from this figure, in comparison to Sourmash, `YACHT` experiences significantly fewer false positives and equivalent false negatives in the uniform distribution case, or slightly increased false negatives in the exponential case. This later case of increased false negatives is due to the exponential distribution causing low abundance genomes to have a coverage in the sample well below the `--min coverage` value of 1, similar to what was seen in Figure 10.

**Fig. 13:**
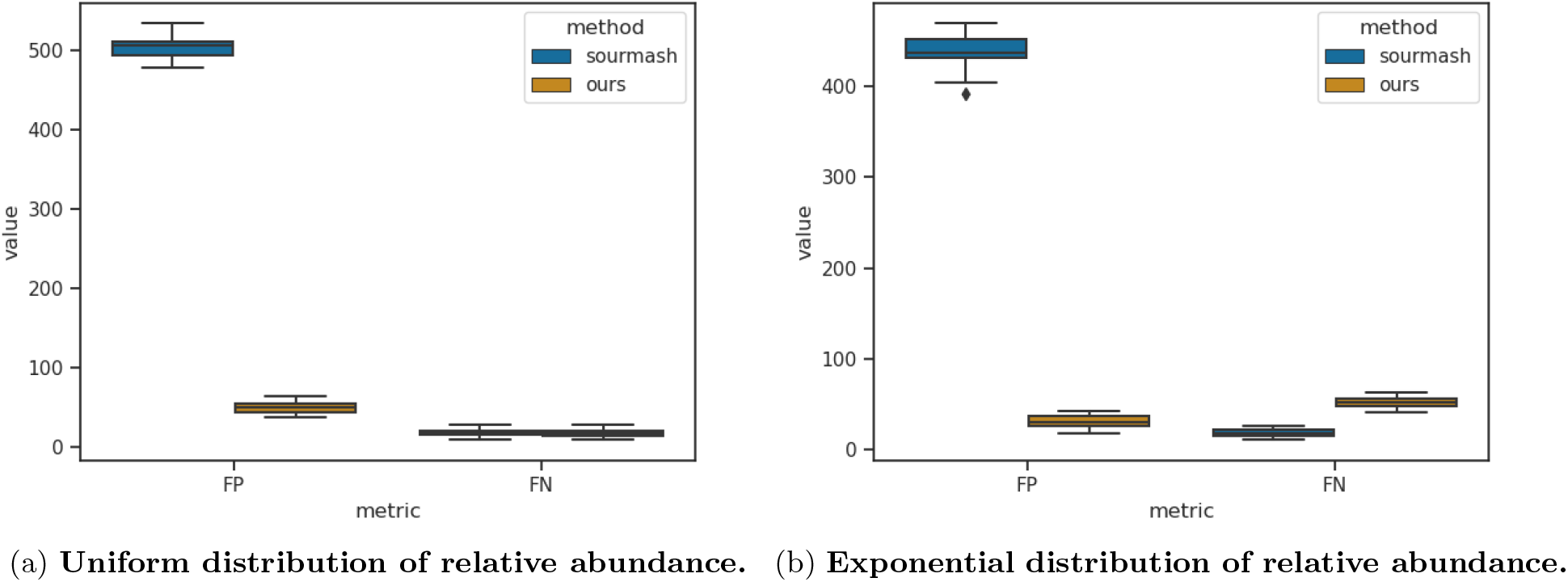
False positive and negative rates of YACHT vs Sourmash. Performance is measured on 30 replicates of simulated metagenomes consisting of 10 million noisy reads generated from 200 randomly selected genomes, half of which were removed from the reference database prior to running Sourmash and `YACHT`. The relative abundance of these 200 genomes was pulled from either (a) a uniform or (b) exponential distribution.

## 6 Conclusion

In this study, we presented `YACHT`, a statistical test for determining the presence or absence of a genome in a metagenomic sample while also accounting for genome relatedness via average nucleotide identity (ANI) and genome coverage in the sample. To our knowledge, this is the first such approach that accounts for these real biological issues while also providing more than just a point estimate for presence/absence, but also an associated confidence. Our synthetic and real-world experiments demonstrate the reliability of `YACHT`, and demonstrate that the theoretical predictions extend to real-world data. Additionally, we showed that this approach is scalable to large data sets due to its reliance on the sketching technique FracMinHash.

We intend this approach to be useful to biologists in distinguishing between low-abundance microbes in a sample and noise, thus facilitating “rare microbiome” analysis with statistical confidence. Additionally, with `YACHT`, it is no longer necessary to apply ad-hoc filtering of low abundance microbes predicted to be in a sample. Rather users can select a significance level (and hence, desired false negative rate), an ANI threshold where they consider organismal genomes as “essentially the same”, and a coverage value by which a genome is judged as being present in the sample. `YACHT` will then perform the filtering via its statistical test.

As such, we envision that `YACHT` can be used as a substitute for Sourmash’s `prefetch` sub-routine. Subsequent application of Sourmash’s `gather` can then be used to associate relative abundances to the genomes deemed present in the sample by `YACHT`.

## Acknowledgements

Alexei Novikov and Stephen White were partially supported by NSF award No. DMS-1813943. David Koslicki and Chunyu Ma were supported by NSF award No. DMS-1664803 and the NIH grant R01GM146462.

## Footnotes

1 More powerful tests exist if one is willing to make distributional assumptions on the ANI *a*, but these will be highly dependent on the underlying biological system.

2 via https://data.gtdb.ecogenomic.org/releases/release207/207.0/genomic_files_reps/gtdb_genomes_reps_r207.tar.gz

